# Accurate Genome-Wide Phasing from IBD Data

**DOI:** 10.1101/2022.04.11.487932

**Authors:** Keith Noto, Luong Ruiz

## Abstract

As genotype databases increase in size, so too do the number of detectable segments of *identity by descent* (IBD): segments of the genome where two individuals share an identical copy of one of their two parental haplotypes, due to shared ancestry. We show that given a large enough genotype database, these segments of IBD collectively overlap entire chromosomes and can be used to phase them accurately. Furthermore, primarily using instances of IBD that span multiple chromosomes, we can accurately phase an entire genome.

We are able to separate the DNA inherited from each parent completely, across the entire genome, with 98% median accuracy in a test set of 30,000 individuals. We estimate the IBD data requirements for accurate phasing, and we propose a method for estimating confidence in the resulting phase. We show that our methods do not require the genotypes of close family, and that they are robust to genotype errors and missing data. In fact, our method can impute missing data accurately and correct genotype errors.

**Author summary:** We present a method for *phasing*, separating the DNA inherited from each parent, of an entire genome using short segments of DNA that are shared between the genome of the person we wish to phase and those of distant cousins. Essentially, if we have enough of these distant cousins’ genotypes available, we can piece together overlapping segments until we have phased the entire genome.

We have developed a method that can phase large numbers of individuals, and that makes special considerations for potential close family and potential genotype errors in data.

We report results on experiments phasing 30,000 individuals. We analyze how many such segments are required to phase accurately, and propose a model-based method to estimate accuracy. We also show that our method can accurately impute missing data and correct genotype errors.

## 1 Introduction

*Phasing* refers to the separation of maternally and paternally inherited DNA. Genotype data are most often generated in an unphased state, because genotyping technologies work at a local level, determining the diploid genotype of one single-nucleotide polymorphism (SNP) at a time. However, phased data are often significantly more valuable. For example, some genotype-phenotype relationships depend on how certain variants are situated across both copies of a homologous genomic region, and phase information is beneficial in the study of genomic diversity and for the purpose of haplotype matching.[5] In applications where pedigrees are available, there is an advantage in knowing which genomic variants of interest are inherited from the same parent by having genomic data phased across the whole genome, and in any application regarding the ancestry of an individual it is advantageous to consider the entire genome, and to consider the DNA inherited from each parent as having its own ancestry.

The advantages of phased data are known and there are several methods of modeling and analyzing genotype data for the purpose of phasing, *e*.*g*., [1, 3, 4, 6] but they are often only able to phase well at a local level–the two parental copies are inevitably swapped many times across the genome.

A simple and common way to separate the haploid DNA inherited from each parent is to compare them to the data of the parents themselves. This is *duo*-phasing, or *trio*-phasing if both parents’ genotype data are available. However, the parents’ data may be expensive or impossible to obtain, and if parental data are missing or all three individuals are heterozygous at a site, the phase cannot be resolved with Mendelian logic alone, and one must defer to a model.

*Identity by descent* (IBD) occurs when one of a person’s two haplotypes is identical to one of another person’s in a segment of the genome because the two share a common ancestor. It has been previously shown that IBD data can be used to phase and determine the parent from which haplotypes are inherited.[2] The approach is essentially to identify segments of IBD and duo-phase those parts. This is the crux of our approach as well, except that in our scenario, there are enough IBD segments that we can expect most IBD segments to overlap others on both sides of the family, and most sites to overlap multiple IBD segments, each providing information on which allele is part of a shared haplotype even if those IBD segments are between unphased diploids. Figure 1 shows a small illustration of the type of data we use.

**Figure 1:**
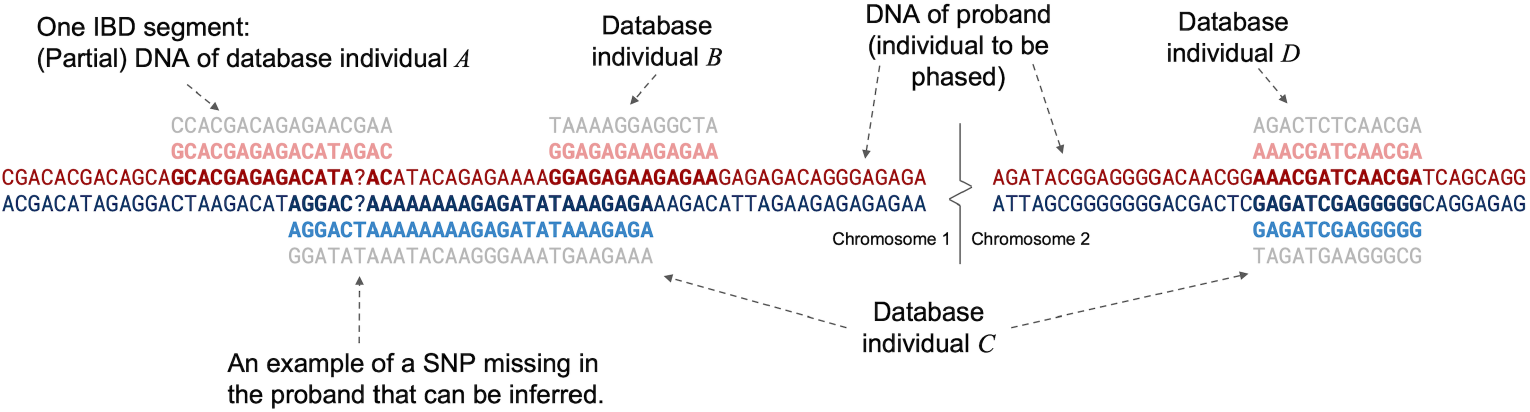
An illustration of the genotype data used for phasing. The DNA of the proband consists of two haploid genotypes across all chromosomes. IBD segments also consist of two haplotypes, one of which is identical to one of the proband’s haplotypes. Note that each IBD segment (partial diploid) is consistent with exactly one of the proband’s haplotypes, and that we can infer that the same haplotype in the proband is shared with individual *A* and individual *B*, even though those segments do not overlap (because they both overlap with individual *C*). Individual *C*’s two IBD segments are on different chromosomes and it is more likely than not that the haplotypes shared with individual *C* are inherited from the same parent (with dozens or hundreds of multi-chromosome IBD segments, inter-chromosome phasing becomes clearer).

When IBD segments overlap, they form a collective block of the genotype for which there is essentially only one way to assign each IBD segment to one parental haplotype or the other, and infer the phase of the proband. Given enough overlapping IBD data, these blocks extend to the full size of a chromosome, although they are not guaranteed to do so. We refer to these blocks as *subclusters*: a block of overlapping IBD segments separated into two parental groups.

The task that remains is to determine which parental group of each subcluster corresponds to the same parent in the other subclusters. That is, we must phase the subclusters into larger *superclusters*, and align the parental groups of each subcluster. Note that we cannot determine from autosomal DNA which parent is the mother. Our goal is to combine all subclusters into one supercluster that separates all the IBD segments into two parental groups, *A* and *B*, without necessarily identifying the parents. We illustrate this task in Figure S1 (in Section S1 below).

The primary mechanism for aligning the parental groups of subclusters on different chromosomes is based on individuals that share IBD segments across multiple subclusters. For example, in Figure 1, individual *C* shares DNA with the proband on chromosomes 1 and 2. It is generally more likely that both the proband and individual *C* inherited both of the shared haplotypes from the same side of the family rather than (*e*.*g*.,) individual *C* sharing DNA with the proband’s mother on chromosome 1 and with the proband’s father on chromosome 2, although the latter case is possible. However, when our IBD data consist of many instances where the proband shares DNA across multiple subclusters, we can phase the subclusters correctly by maximizing the number of instances where multi-segment IBD data are inherited from the same parent in a supercluster.

Depending on the amount and distribution of IBD data, it is possible that some parts of the genome do not overlap any IBD data and cannot be phased (in which case we may default to the phase inferred by models), or that some subclusters cannot be connected to others through multi-segment IBD (in which case we will have multiple superclusters of varying size), but we evaluate our method on the genome-wide phase that results in all scenarios and characterize the IBD data required to apply our approach. We use this method to phase 30,000 child-father-mother trios using IBD segments detected among a database of 12,755,111, excluding IBD shared with parents. We call our proposed method **IBDphase**. In the Sections that follow, we show the phase accuracy of the method and that this approach can be used to effectively impute missing data and correct genotype errors in data. We provide details of our approach, including how we make our methods robust to potential genotype errors in the proband genotypes, and propose a method for estimating the accuracy of the resulting phase.

## 2 Results

The primary methodology we use to evaluate our approach is to use the genotype data of 30,000 child-parent-parent trios, identify IBD shared between the children in this test set and a database of over 12 million genotypes, apply IBDphase without using the parents’ genotype or IBD data, and then compare the resulting phase to trio phase using the parents’ data. The statistic we are most interested in is the global separation of the alleles inherited from each parent: when we phase the genome into two haploid sequences, to what extent did we correctly separate all of the alleles inherited from mother from all the alleles inherited from father? We also measure local phase accuracy, and the accuracy of the assignment of IBD segments to parental sides. We describe our experimental methodology in detail in Section 4.3.

We pre-phase our test set using Eagle v2.4.1[6] (using only the cohort of 30,000 individuals). We do not consider the resulting phase accuracy to be the state-of-the-art standard for comparison, but pre-phasing our data is a necessary step for our approach because it provides a default to make phasing decisions at sites where there are no overlapping IBD segments. Indeed, methods like Eagle are not suited to separate the DNA inherited from each parent at a genome-wide level. In contrast, IBDphase is able to separate the DNA inherited from each parent in our test set with an average accuracy over 95% (and a median accuracy of nearly 98%). Phase accuracy results are shown in Table 1 (see also Table S1 and Figures S5 and S6 for complete statistics). IBDphase also provides excellent local phase accuracy in these experiments. The median resulting “phase flip” rate is 0.26%, and most of the test set have at least 99% of their heterozygous SNPs phased in complete agreement with trio phase in a region that measures at least one centimorgan (this is a measure of local phase accuracy that does not penalize for small areas of the genome that are phased very poorly).

**Table 1:** Phase Accuracy. The genome-wide phase accuracy of IBDphase. Accuracy is measured using trio phase as the standard of true phase, using only SNPs where the phase can be unambiguously inferred from the trio (*i*.*e*., where at least one parent is homozygous). Global error is the rate at which the phase differs from trio phase (keeping only one haplotype assigned to one parent across the genome, but assuming the more favorable haplotype of two choices). Phase flip rate is the frequency with which the phase of a heterozygous SNP differs from that of the previous heterozygous SNP with respect to trio phase. The third accuracy measure is the proportion of SNPs that belong to segments where there are no phase flips for at least 1 centimorgan. Some measures depend on SNP density, and we consider 416,176 SNPs across the autosome. The median global phase error of the pre-phased data is 48%, and the median phase flip rate and median proportion of SNPs in 1 cM+ runs is 1.08% and 95% respectively.

| Criteria | IBDphase median | IBDphase mean |
| --- | --- | --- |
| Global phase error | 2.09% | 4.93% |
| Phase flip rate | 0.26% | 0.33% |
| Proportion of SNPs in 1 cM+ runs | 99.01% | 98.64% |

IBDphase also labels each match segment as being on one side of the family or the other. When we compare the groups created to the IBD segments we identify using the parents’ DNA, we find that the average IBD segment assignment error is 3.4% and the median error is 0.67%. This measure of error is comparable and correlated with genome-wide phasing error (See Figure S11), but does not penalize for portions of the genome that do not overlap IBD segments.

### 2.1 Estimating the Accuracy of Resulting Phase

IBDphase performs better when the genomic database is large, when many IBD segments are discovered in it, when a large proportion of sites overlap at least a few IBD segments, and when there are close genetic relationships to provide long IBD segments and help phase across multiple chromosomes. Table 2 shows that all these measures are strongly correlated, and provides an estimate of how performance on this database may continue to improve. As expected, the global error rate at sites that do not overlap IBD is high (median value 30.5%) as is the overall phase error when the database is too small. (See Table S1 and Figs. S4, S5, S6, and S7 for more information, and Supplementary Data File 1 for detailed data.)

**Table 2:** Performance and IBD statistics as database size increases. As the size of the genomic database in which IBD is identified increases, so does the global phase accuracy, number of close genetic relationships, proportion of the genome that overlaps at least a few IBD segments, and the overall number of IBD relationships.

| Database size (Millions) | Median Phase Error | Median number of close (400 cM) relations | Median 5x overlap rate | Median number of IBD relations |
| --- | --- | --- | --- | --- |
| 1.00 | 36.8% | 1 (mean 1.32) | 67.7% | 5,024 |
| 2.00 | 22.1% | 1 (mean 1.58) | 82.7% | 10,003 |
| 5.00 | 6.99% | 2 (mean 2.43) | 94.9% | 24,623 |
| 12.76 | 2.09% | 3 (mean 4.31) | 98.7% | 60,778 |

The correlation between these statistics and observed phase accuracy at the level of an individual proband is much weaker, however (*e*.*g*., see Fig. S7), but we can combine features of the phasing process to train a predictor that can help estimate the confidence we have in the phase quality of any individual proband (see the Methods section for details). Table 3 shows the most informative features. The Pearson correlation between the prediction from our error model on this test set and the observed global phase error is 0.66 and the Spearman correlation is 0.45.

**Table 3:** Features used to estimate phase confidence. The top 10 features, ranked by their importance (GINI importance, as calculated by *scikit-learn*[11] for random forests).

| Feature | Importance |
| --- | --- |
| Proportion of database individuals with IBD segments assigned to both sides of the family, both on the largest supercluster | 0.388 |
| Proportion of IBD segments that are partially assigned to one parental side and partially to the other | 0.108 |
| Number of close family members that do not share DNA with all other close family | 0.103 |
| (log) number of database individuals with shared DNA | 0.088 |
| (log) number of IBD segments assigned to the largest supercluster | 0.081 |
| Number of close family members that do not share DNA with all other close family | 0.032 |
| Proportion of the genome overlapped by at least one IBD segment | 0.030 |
| Ratio of the number of database individuals with shared DNA on one IBD segment to the number with shared DNA on multiple IBD segments | 0.029 |
| Proportion of the genome overlapped by at least two IBD segments | 0.021 |
| Proportion of IBD segments assigned to the largest supercluster | 0.018 |

A large proportion of our database consists of individuals with European ancestry and as expected, the performance of IBDphase is worse on individuals whose ancestry is underrepresented in our database. There is strong evidence that the decreased performance in such individuals is due to the fact that there is a smaller number of IBD segments associated with them. For example, if we classify each test set individual according to the major geographic region that explains the largest portion of genetic ethnicity according to AncestryDNA estimates (there are 12 such regions with at least 20 representative individuals, see Table S2 and Supplementary Data File 1), we observe that the proportion of the genome that is overlapped by at least five IBD segments is a much stronger predictor of phase accuracy than the genetic ethnicity designation. Specifically, if we divide the amount of genome coverage into 12 exclusive ranges, the mutual information between genome coverage and the decile of the phase error is 0.24, whereas the mutual information between the genetic ethnicity designation and the phase error is 0.027. Indeed, if we restrict our evaluation to individuals with a minimum genome coverage, the performance improves across all groups (Table S2). We also observe that the performance of IBDphase in admixed individuals such as African Americans (in this case, defined by having at least 20% African and 20% European estimated assignment and at least 80% combined African and European assignment) is comparable to the rest of the test set (median global phase error 2.60%, n=1587), and the median global phase error in Latinx test set individuals (10% European, 10% from the Americas, 50% combined European and American) is 2.09% (n=1039). These observations suggest that as the database continues to grow, the performance of IBDphase increases, regardless of demography.

### 2.2 Imputation and Correction of Genotype Errors

IBDphase can also improve imputation accuracy and correct genotype errors because it observes the genotypes of several individuals at the same site (the proband and overlapping IBD segments). In cases where overlapping IBD segments imply a different genotype than the one in the input or imputed in the pre-phased data, IBDphase will override the call. We test the effectiveness of these corrections by artificially altering some genotypes and setting some genotypes to missing before performing the same phasing experiment as above on a random 5,000 of our original 30,000-individual test set. In these experiments, we evaluate IBDphase by whether or not it is able to replace the perturbed genotypes with the original. Since artificial genotype errors affect whether or not IBD is detected, we evaluate only on SNPs that are not used in IBD detection (although results are similar when we evaluate on all SNPs–see Supplementary Data File 1).

In the scenario where we set a genotype to missing, a genotype is imputed during the pre-phasing process (again, we use Eagle[6], this time on the 5,000 individuals whose data we perturbed and the 25,000 unaltered members of the original test set). When there are not enough overlapping IBD segments on either side of the family, IBDphase always keeps that imputed call, which is correct 97% of the time. When there is enough overlapping IBD on one side only, we change the imputed call 1% of the time, and the change is correct 97% of the time. When there is enough IBD on both sides, we change the call 0.5% of the time, and the change is correct 95% of the time. In this experiment, IBDphase reduced the number of imputation errors from 86,935 to 40,689 (a 53% reduction).

In the scenario where we replace a genotype with a genotype error, the pre-phased data always reflects the perturbed genotype, and if there is not enough overlapping IBD on either side of the family, IBDphase keeps that call, which is always incorrect in this simulation. However, when there is enough overlapping IBD on one side of the family, IBDphase overrides the call 50% of the time, and the change is correct 100% of the time (with IBD on one side, IBDphase only overrides homozygous calls). When there is enough overlapping IBD on both sides, IBDphase overrides the call 79% of the time and is correct 99.993% of those. IBDphase corrected 65% of all genotype errors in total. When the given genotype was not altered for this experiment, IBDphase overrode the call in 0.017% of the instances when there was enough overlapping IBD on one side, and 0.020% of the instances when there was enough IBD on both sides. Accounting for all SNPs in this experiment, perturbed or not, IBDphase reduced the number of genotype errors by 55%.

## 3 Discussion

IBDphase provides excellent local and genome-wide phasing, as well as the assignment of IBD segments to parental haplotypes, provided a large enough IBD database. Our method is able to phase, impute, and correct errors with high accuracy at a local genomic level, and accurately identify the corresponding parents in phased data across chromosomes, resulting in complete separation across the genome of the DNA inherited from each parent, with an accuracy usually above 95% (accuracy is over 95% more than 75% of the time, and above 99% over 30% of the time). IBDphase also labels IBD segments in a cluster hierarchy that indicates the genomic locations of the most confident resulting phase, and provides an error model that can predict with reasonable accuracy whether or not the phase is likely to be reliable genome-wide. Our approach is comparable to the techniques used in Kong *et al*. [2] but our aim is to separate maternally and paternally inherited DNA across the entire genome without requiring an elevated level of IBD sharing in any specific population, as the deCODE study has in Iceland, which requires that database to be substantially larger. We measure the accuracy of genome-wide phasing relative to the size of a database and the distribution of IBD segments we can identify within it.

After IBD segments are separated into parental groups, breakpoints become much more apparent, where numerous IBD segments end and another set of IBD segments begin, which are likely to be places where a recombination event occurred in one of the proband’s ancestors. One area for further study is the extent to which these groups can be associated with specific ancestors and how large a database is necessary.

One of the potential pitfalls with our method is an instance where a significant number of individuals share DNA with both of the proband’s parents. These are often cases where the proband’s parents are not closely related, but do have ancestry that is similar enough that they share DNA with the same individuals due to identity by state or founder effects, or from influences of SNP density and population-specific allele frequencies. Such scenarios add noise to the information IBDphase uses to phase subclusters, as evidenced by the fact the most informative feature in estimating the phase quality is the number of database individuals IBDphase is forced to put on opposite sides of the family (Table 3). Distinguishing IBD segments that are not informative in this way from those that are informative is a difficult open problem and effective solutions are likely to provide the largest benefit to the performance of IBDphase.

## 4 Methods

### 4.1 Our IBD-Phase Approach

Our approach has the following steps (for each proband we wish to phase)

1. Load the IBD coordinates (specifying which individuals in the database share IBD with the proband), as well as the genotype data for those individuals at those coordinates, all genotype data for the proband, and all genotype data for close family of the proband (if any). If available, load phased data for the proband (these data will provide the “default” phase for sites that cannot be resolved from IBD segments. If not available, the default phase is arbitrary).
2. Compare the genotypes of close family to each other and determine if they are potentially other descendants of the proband’s parents (we will discard those).
3. Identify subclusters: groups of IBD segments such that each significantly overlaps with others in the same group.
4. Separate the IBD segments of each subcluster into two parental groups, potentially breaking some into smaller pieces where data are inconsistent.
5. Phase the subclusters and group them into superclusters and determine which parental group of each subcluster corresponds to which parental group of the supercluster.
6. Phase the proband genotype across the genome using the genotype information in the separated parental groups.

We describe each of these steps in turn. First, we load data. For efficiency, it is essential that genotype data are stored in binary format, indexed by individual, and all use the same coordinates (*i*.*e*., set of SNPs) as the IBD data. Then, for all IBD segments, we can seek and read only the segments of genotype data that correspond to the IBD segments.

Second, we compare the genotypes of close family. A close family member is anyone who shares a significant amount of IBD (*e*.*g*., 400 centimorgans) so that we can safely assume that if they are related to both of the proband’s parents, they will also be related to other close family. Individuals who are descendants of the proband’s parents (*e*.*g*., a nephew) are unreliable data for phasing the proband, because they will share DNA with the proband throughout the genome on both sides of the family. We discard any IBD segments from close family that shares IBD with all other close family because they are potentially descendants of both the proband’s parents. We also discard IBD segments shared with an identical twin or full sibling of the proband (which are relatively easy to identify as such). It is possible that we may discard IBD shared on only one side of the proband’s family. In contrast, close family that are not discarded are particularly useful in phasing the proband because they will share a significant amount of DNA with the proband exclusively on one side of the family. Note that close family are not required for our approach, but if they are available we may label them and prefer to use them in cases where phase information conflicts with that of other IBD segments.

The next step in our approach is to delineate the IBD segments that make up subclusters. These are collections of IBD segments that overlap on the genome enough to determine whether they are on the same side of the family or the opposite. That determination is primarily based on homozygous sites in IBD segments because there is only one allele which therefore must be the allele shared with the proband. We cannot assume that our IBD data perfectly delimit where a shared haplotype begins and ends. Such IBD data are typically generated by comparing pairs of diploid genotypes in unphased or imperfectly phased data, and IBD is commonly estimated as extending beyond the genomic positions where the identity is due to a shared ancestor. We therefore insist that each IBD segment in a subcluster overlaps another in the same subcluster by a minimum number of sites (*e*.*g*., 40) that are heterozygous in the proband but homozygous in both IBD segments so that we are confident that each IBD segment will be placed into parental groups correctly with respect to the other IBD segments. Under this constraint, it is straightforward to build subclusters and each will overlap part or all of a chromosome.

The next step is to separate the IBD segments of each subcluster into two parental groups (an illustration of our approach is shown in Figure S2). To make this efficient, we rely on the fact that (with very few exceptions which are discussed below) any two overlapping IBD segments will either share the (i) same or (ii) the opposite haplotype with the proband, and will therefore (i) always be homozygous for the same allele or (ii) always be homozygous for the opposite allele at sites where the proband is heterozygous and both IBD segments are homozygous. For example, in Figure 1, in all the places where the proband is heterozygous but two IBD segments are both homozygous at the same site, the two IBD segments are homozygous for the opposite alleles, implying that they do not share the same haplotype. Our approach is to scan the proband’s heterozygous sites and divide the overlapping IBD segments into two groups: those that are homozygous for one allele, and those that are homozygous for the other. These represent the two sides of the proband’s family, and our aim is to make sure that all IBD segments are consistently assigned to the same group. In practice, there are conflicts to resolve which arise either from genotype errors, or from the aforementioned fact that several IBD segments are likely to extend beyond the genomic positions where they truly share a haplotype. We account for this by downweighting the information contribution of an IBD segment if a SNP site in question is near either end of the segment (see Fig. S9). At each site that is heterozygous in the proband, we choose to phase the site in whichever way maximizes the (weighted) number of IBD segments that will stay in the same parental group to which they were assigned at the previous heterozygous site in the proband. If there are any IBD segments that must change sides of the proband’s family as a result, we break that segment into two. In practice, few segments are significantly broken and we are left with an improved estimate of their positions (See Table S3 for relevant statistics from our experiments). There is one potential pitfall that we must account for explicitly: if the proband is *erroneously* called heterozygous, then the procedure described above is likely to break up segments unnecessarily and assign them to the wrong parental groups. It is not always clear when examining individual SNPs that the proband is homozygous but called in error, and once the parental groups of IBD segments have become corrupted as a result, they can cause further errors downstream of the erroneous site. To address this issue, we employ a lookahead scheme. We associate a cost with breaking up an IBD segment (*i*.*e*., changing its assignment from one parental group to the other), and a cost with ignoring a site. If the cost we incur at a site that was called heterozygous in the proband and several other heterozygous sites both upstream and downstream of it exceeds the cost of ignoring the site, then we leave the parental groups intact. In our experiments, we ignore about 0.3% of SNPs. Note that if an IBD segment shares *both* haplotypes (called “IBD2”), it will not affect the phase. All sites that are heterozygous in the proband will be heterozygous in the other genome and the segment will not be assigned to either side of the family. Only in cases where an IBD2 segment has several genotyping errors and no other IBD1 segments to “outvote” it will IBD2 data infer incorrect phase. Full siblings are the only relationships with significant IBD2, and these are straightforward to identify and discard.

Once the subclusters are defined, and all IBD segments within them are separated into two groups, the next step is to align the subclusters’ parental groups and combine them into larger “superclusters”. If the proband shares DNA with the same database individual on two different IBD segments in two different subclusters, we prefer to build a supercluster that puts both of these IBD segments in the same parental group. Our objective is to maximize the number of such instances, possibly weighting them by the amount of DNA shared with the proband. We are not guaranteed to have enough multi-segment IBD instances to combine all subclusters into exactly one supercluster, but in practice, the majority of IBD segments belong to the largest supercluster (96% on average, in our experiments) In fact, if our purpose is to cluster IBD segments by parent or if we wish to delimit the parts of the genome with the highest confidence global phase, we may choose to insist on a minimum number of multi-segment IBD instances to connect subclusters together. We choose a simple greedy optimization approach with random restarts to align the parental groups of subclusters which is efficient and works well in practice (See Figure S1). We begin by grouping subclusters into superclusters such that all subclusters have a minimum amount of multi-segment IBD to connect to the larger supercluster. For each supercluster, we begin by phasing the subclusters randomly (*i*.*e*., align subcluster parent *A* randomly with supercluster parent *A* or *B*), then greedily swapping the assignments of whichever subcluster increases the objective score the most. We repeat the random-initialization and greedy optimization some number of iterations and keep the best subcluster phase.

Once IBD segments are assigned to superclusters and their parental groups, we phase the proband across the entire genome. We choose to do so at this point because (i) we have now resolved the phase of potentially overlapping subclusters, and (ii) we are interested in examining not just the heterozygous sites in the proband, but also imputing missing data and potentially overriding the original calls if there is enough evidence in overlapping IBD segments to do so. Because the IBD segments are organized into two parental groups, the phased genotype of the proband can be inferred from the homozygous sites in those groups’ overlapping IBD segments. In this step of our approach, we have the option to insist on a minimum number of overlapping segments in order to override the original calls, and we may also make use of a pre-phased version of the proband (*e*.*g*., inferred from phasing models) to make inferences at any site that does not have enough overlapping IBD segments.

### 4.2 Estimating Phase Confidence

A minority of individuals in our test set are phased inaccurately, with genome-wide phase accuracy as low as 50% (*i*.*e*., the alleles inherited from each parent are evenly mixed throughout the genome). These individuals may have accurate local phase, but our goal is to separate the DNA inherited from each parent on a genome-wide scale. The genome-wide phase accuracy depends on the ability of IBDphase to identify the DNA inherited from the same parent in phased data (our subclusters) across different chromosomes and that accuracy is not a function of the number or distribution of IBD segments that applies in all cases.

To identify low-performing cases, we build a random forest regression model that uses the following features.

- Close family:the number of immediate family (generally second cousin or closer relationships) that can be identified as being on only one side of the family.
- IBD statistics: number of database individuals with whom IBD is identified, proportion of those with multiple IBD segments, and proportion of those with exactly one segment.
- Clustering features: number of IBD segments linked to the largest supercluster, number of database individuals assigned to opposite parents within that supercluster, mean and median ratio between the majority and minority parent side that IBD segments are assigned to.
- IBD genome coverage: the proportion of the genome that overlaps a minimum number of IBD segments. One feature for each of 1x, 2x, 5x, 10x, and 20x coverage.

We train the error model on a separate set of 25,641 trios (*i*.*e*., these are not the 30,000 individuals used to generate the results above), and use a set of 6,410 trios for validation. We train our model using *scikit-learn*[11] with a parameter grid to optimize the model, which we use to predict the accuracy of the parent assignment to IBD segments in the largest supercluster in the validation set. The relationship between predicted and observed phase accuracy on the test set of 30,000 is shown in Figure S8.

In practice, we may use the estimated error to make more conservative judgements about our IBD segment clusters. For example, if the predicted error is over 20% (which is the case for about 5% of individuals), we replace the largest supercluster with the largest *sub*cluster, expressing a higher degree of confidence on a smaller portion of the genome, before calculating the accuracy in Figs. S6 and S11.

### 4.3 Experimental Methodology

We use the following procedure to carry out all of our experiments. We identify 30,000 parent-parent-child trios (90,000 unique individuals) from our database by identifying triples such that the first individual shares DNA on one haplotype across approximately 100% of the autosome with the second and third individuals, but the second and third individuals do not share a substantial amount of DNA with each other (the parents may share up to approximately 400cM, which still makes parent-parent-child the only feasible relationship scenario).

We identify IBD shared between members of this test set and a database of approximately 12.7 million individuals. There are several available methods for identifying IBD[8, 10, 9, 7] but for the purposes of our experiments we define an IBD segment to be any section of a chromosome of significant length such that two individuals share at least one allele throughout the segment (*i*.*e*., the pair do not have a homozygous opposite genotype for any SNP in the segment). We consider an IBD segment to be of significant length if (i) it is at least 8 centimorgans, and (ii) it measures least 5 centimorgans when we discard any recombination distance between two adjacent SNPs that exceeds 0.05 centimorgans (the second criterion is designed to ensure that any IBD segment we use is supported by a minimum SNP density). We identify these IBD segments using unphased data, which has the advantage of not relying on any properties of phasing quality at all, and we can do this efficiently by first organizing all genotype data into bitmaps that represent the SNPs where individuals are homozygous for either allele, then comparing the genotypes of pairs of individuals over several SNPs simultaneously using bitwise arithmetic. The basis for our procedure is illustrated in Figure S10.

Before we run IBDphase, we generate a “pre-phased” version of each trio child in our test set that will be the proband in the experiments. We emphasize that this pre-phasing step does not affect the assignment of IBD segments to either side of the family, and only influences the phased results in positions where IBD segments do not have the homozygous genotype data that determine the phase. However, without such a phased version to default to, each such site would be phased no better than random guessing. For this step, we use Eagle v2.4.1[6] and provide only the cohort of 30,000 test set trio child individuals.

We provide the pre-phased version of each trio child, the unphased genotype database (including those of the test set), and the IBD segments for each trio child (with those segments shared with parents removed from the list) to IBDphase, which produces the genome-wide phased version of each test set child, as well as the subcluster, supercluster, and parental side labels for each IBD segment. Subclusters consist of IBD segment groups such that each segment in a subcluster overlaps with another segment in the same subcluster by at least 40 sites that are heterozygous in the proband and homozygous in both IBD segments. IBDphase will break up segments if different parts of them are assigned to different parents, but the longest portion of all IBD segments is always retained for the purpose of evaluating their parental side assignment, and all segment portions greater than 5 centimorgans are retained (although if two such segment portions are separated by only a single SNP, they will be considered one segment). To evaluate the results, we introduce the genotypes of the parents in our test set and measure the proportion of heterozygous genotypes in the trio children that are phased in agreement with the genotypes of the parents. Note that the genome-wide phase that results from IBDphase has its alleles divided into two genome-wide haplotypes, but does not associate either specifically with the proband’s mother or father. We consider the genome-wide phase to be accurate if haplotype one agrees with the genotypes of the mother and haplotype two with the father, or if haplotype two agrees with the mother. In this step, we consider all SNPs where the correct phase can be inferred from the parents’ genotypes (*i*.*e*., at least one parent is homozygous) and consider the error to be the number of alleles assigned incorrectly allowing for either haplotype-parent assignment, whichever is the better agreement. We also assign a parent side to each IBD segment and compare those to the IBD segments of the parents themselves (See Figure S11). Unlike global phase accuracy, this measure cannot penalize for parts of the genome that are incorrectly phased but have no IBD segments to inform those sites.

For the imputation and error correction experiments, we perturb the genotypes of the test set trio children before the IBD detection step by randomly replacing 1% of the calls with missing data and changing the genotypes for 0.2% of the calls either from heterozygous to homozygous or from homozygous to heterozygous. In our experiments, IBDphase infers the genotype of the proband from those of overlapping IBD segments as long as there is at least one overlapping segment (after downweighting the SNPs near the segment’s endpoints by the sigmoid-shaped weighting function described above and shown in Fig. S9) and overrides the imputed call or phase decision given by the pre-phased data if the count of weighted IBD segments exceeds 0.1. Since our IBD detection procedure will not report shared segments that have a homozygous opposite, in these experiments, we detect IBD using a subset of SNPs, and observe whether IBDphase imputed or replaced calls correctly on the SNPs that were not used in IBD detection, although the results are almost identical using all SNPs (see Supplementary Data File 1).

## Supporting information

Supplementary Data File 1 (gzipped Excel .xls format)

## 5 Ethics approval and consent to participate

All data for this research project were from subjects who provided prior informed written consent to participate in AncestryDNA’s Human Diversity Project, as reviewed and approved by our external institutional review board, Advarra (formerly Quorum). All data were de-identified prior to use.

## S1 Supporting information

**Figure S1:**
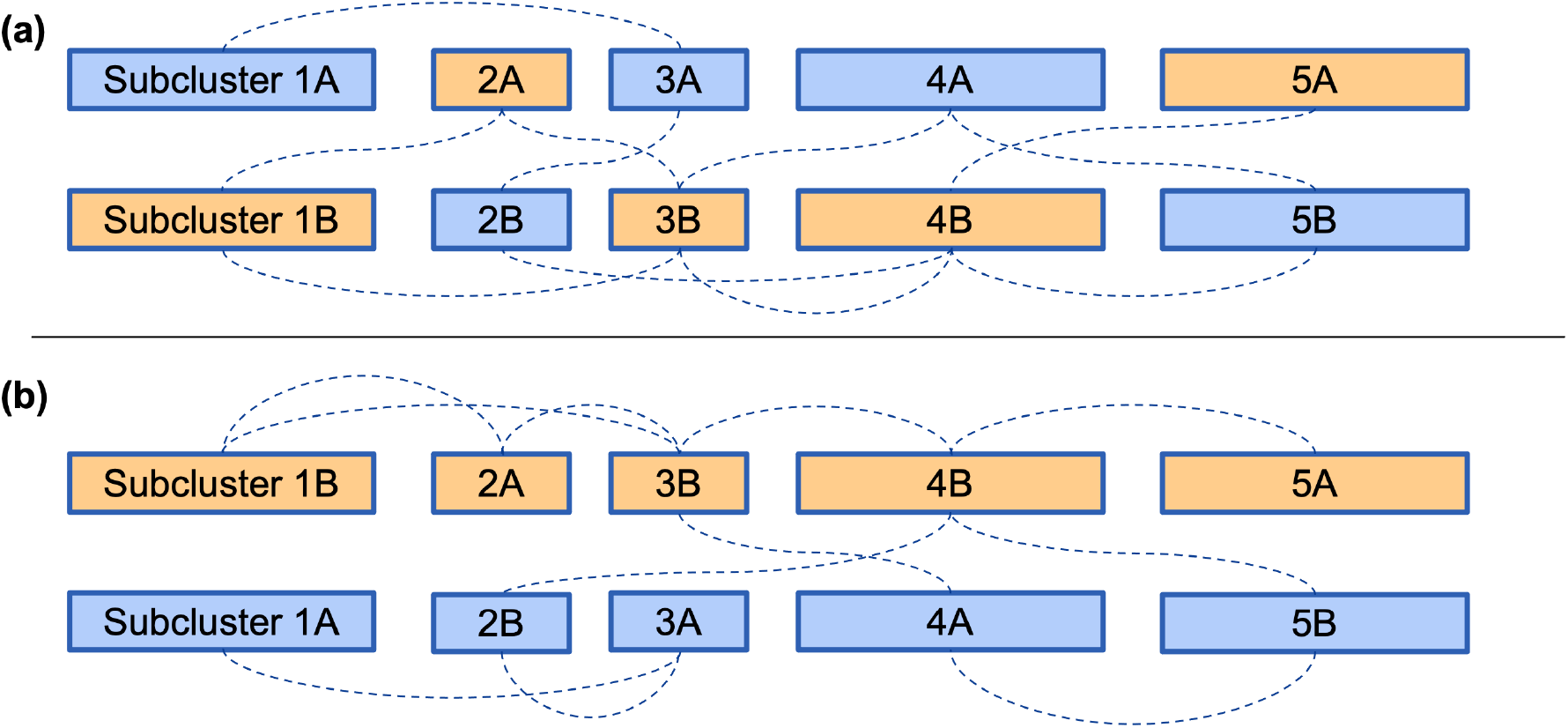
An illustration of the subcluster phasing task. **(a)** shows five subclusters across four chromosomes. The color indicates the true parent from which IBD segments are inherited. Dotted lines indicate a connection between groups where the proband shares DNA with the same person in both groups. **(b)** shows the result of phasing these subclusters into one supercluster such that the number of connections between groups on the same side of the family is maximized.

**Figure S2:**
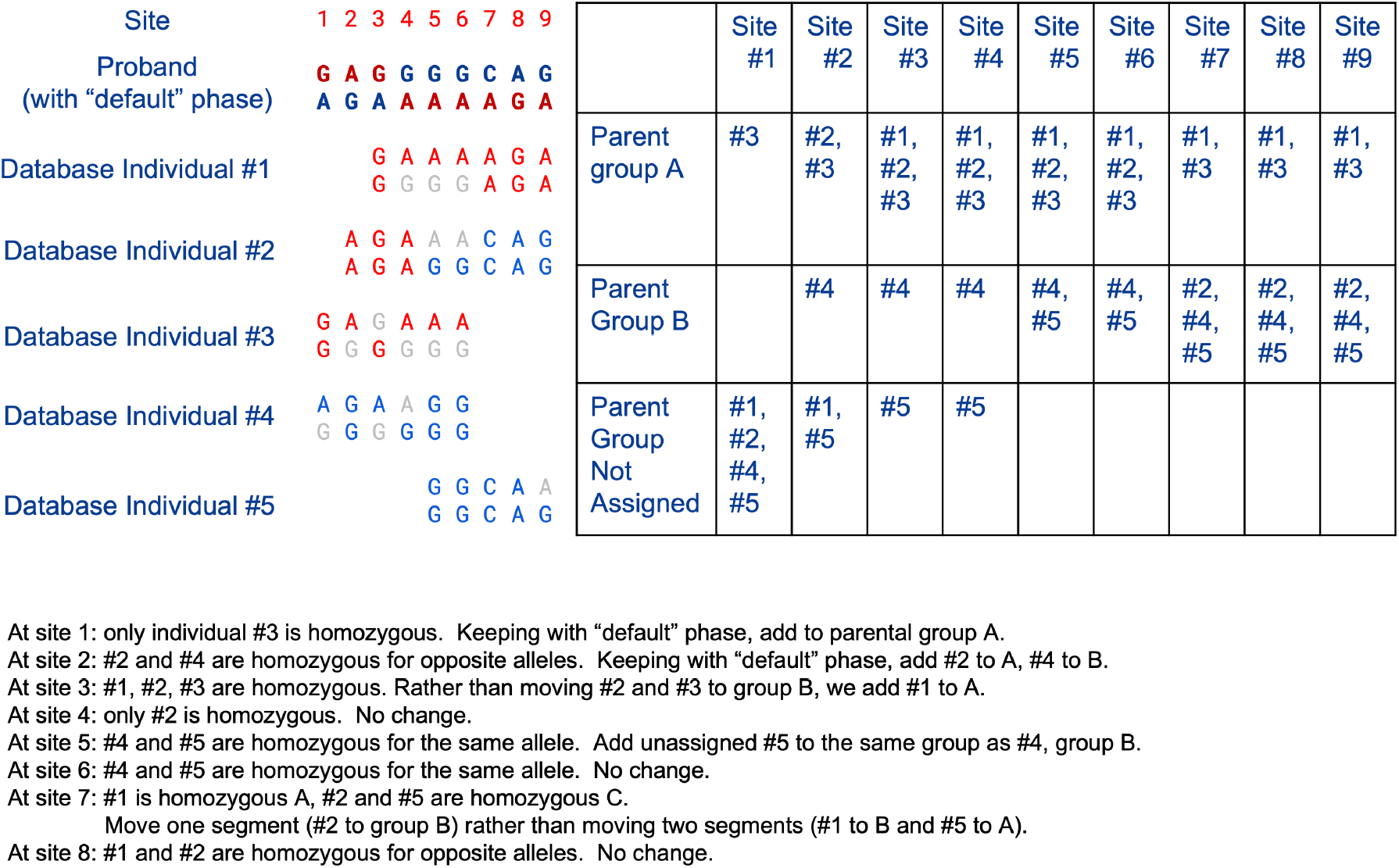
An illustration of the IBD segment assignment task. This toy example shows nine sites where the proband is heterozygous, the “default” phase (*e*.*g*., provided by a phasing model), and five IBD segments. Moving from left to right, we can group the IBD segments into two parental groups based on their homozygous sites: segments that are homozygous for the same allele belong to the same parental group. In this example, there is no way to consistently assign all segments to one group or the other (at site 7, individual #2 who was previously grouped with individual #1 is transferred from parental group *A* to *B*).

**Figure S3:**
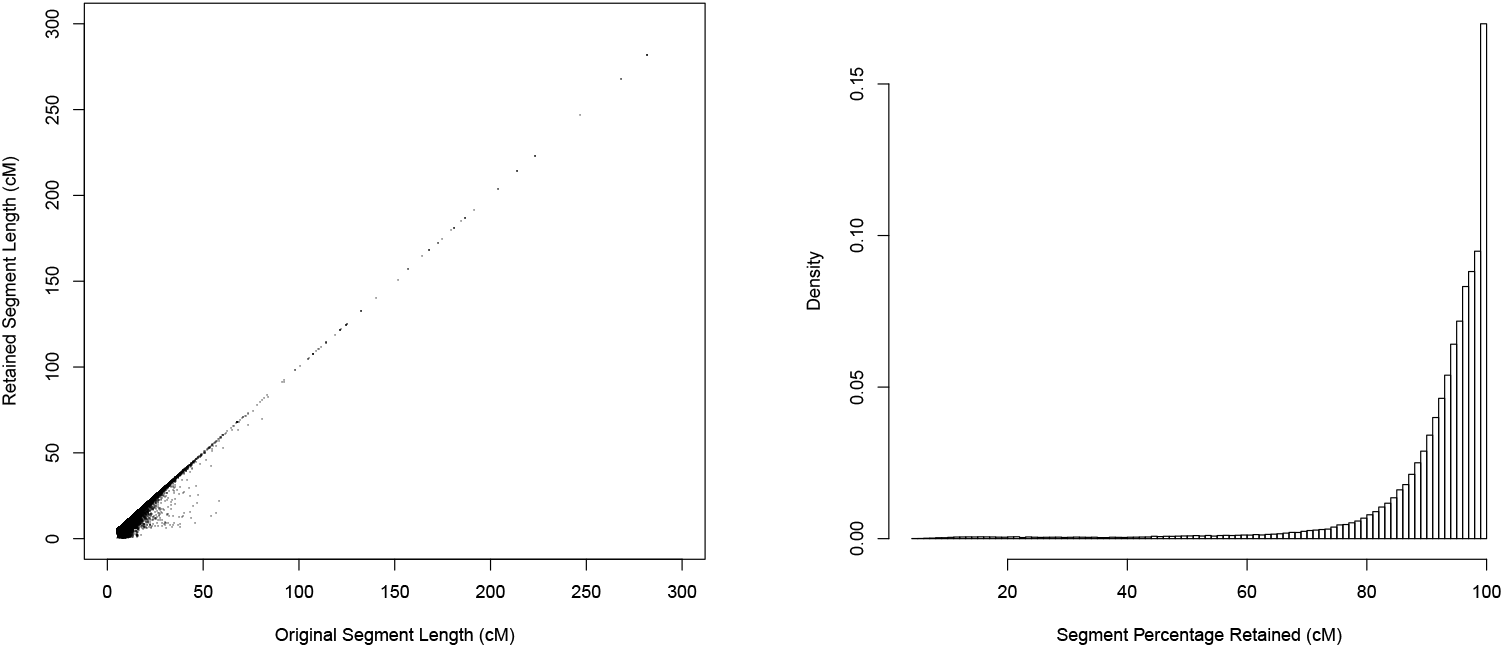
Proportion of segment length retained after assigning parental groups. Both plots show the same data indicating how much of an original IBD segment’s length is retained after the process of assigning each to a parental group (the sample mean is 92% and the median is 96%.) To resolve Mendelian confilicts, some segments must be broken.

**Figure S4:**
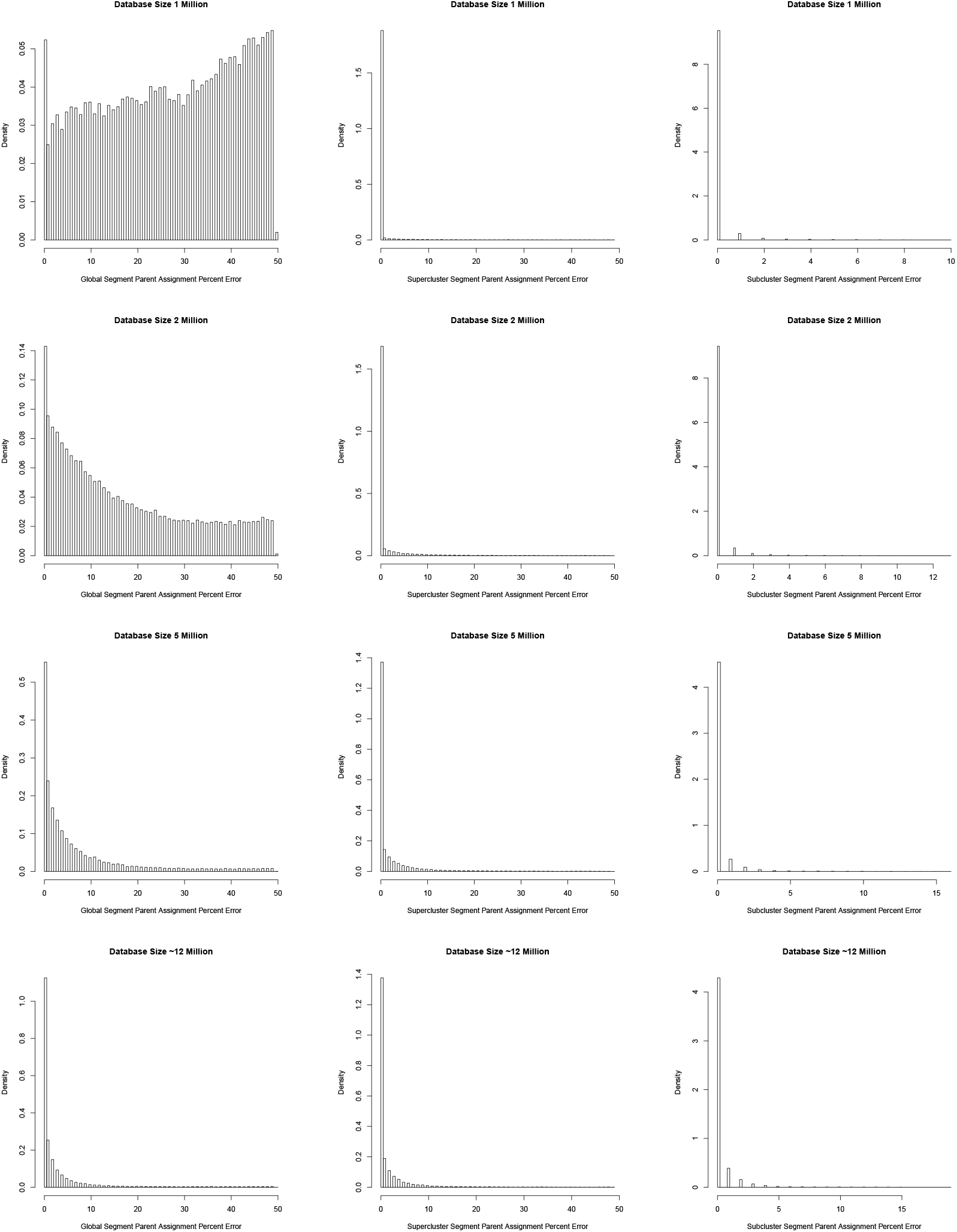
Distribution of phase error based on the size of the database against which we search for IBD. Column 1 shows the global phase error. Column 2 shows the aggregate phase error when each supercluster is evaluated independently. *I*.*e*., if two superclusters both separate maternal and paternal alleles well, they are both evaluated with low error, even if they disagree on which parent is which. Column 3 shows the aggregate phase error when each subcluster is evaluated independently.

**Table S1:** Phase accuracy statistics. Accuracy of phase when (i) Pre-phased (*i*.*e*., using Eagle alone on the 30,000-member test set), (ii) using IBDphase, (iii) using IBDphase, but limited to SNPs in the largest supercluster, (iv) using IBDphase, but limited to SNPs that are covered by at least one IBD segment, (v) using IBDphase, but limited to SNPs that are both in the largest supercluster and covered by at least one IBD segment. The accuracy is showed as measure by (**a**.) Global phase error rate: what proportion of heterozygous SNPs are phased in a way that agrees with trio phase (only counting SNPs that are unambiguously phased by the parents–*i*.*e*., when at least one parent is homozygous), (**b**.) Phase flip rate: what proportion of heterozygous SNPs are phased opposite the previous heterozygous SNP, relative to the trio phase? (**c**.) Average length (cM) of runs of correct phase: what is the average number of centimorgans before a phase flip is encountered? (**d**.) Proportion of SNPs in 1 cM+ runs of correct phase: how many SNPs are in runs of correct phase that are at least 1 centimorgan?

| a. Global phase error rate | Mean | Median |
| --- | --- | --- |
| Pre-Phased | 0.4810 | 0.4842 |
| IBDPhase | 0.0493 | 0.0209 |
| IBDPhase (largest supercluster) | 0.0493 | 0.0209 |
| IBDphase (SNPs covered by IBD segments) | 0.0472 | 0.0195 |
| IBDphase (largest supercluster, covered by IBD segments) | 0.0302 | 0.0118 |
| b. Phase flip rate | Mean | Median |
| Pre-Phased | 0.0124 | 0.0108 |
| IBDPhase | 0.0033 | 0.0026 |
| IBDPhase (largest supercluster) | 0.0033 | 0.0026 |
| IBDphase (SNPs covered by IBD segments) | 0.0032 | 0.0026 |
| IBDphase (largest supercluster, covered by IBD segments) | 0.0027 | 0.0023 |
| c. Average length (cM) of runs of correct phase | Mean | Median |
| Pre-Phased | 2.8057 | 2.8743 |
| IBDPhase | 13.4194 | 11.2563 |
| IBDPhase (largest supercluster) | 13.4194 | 11.2563 |
| IBDphase (SNPs covered by IBD segments) | 13.6185 | 11.4408 |
| IBDphase (largest supercluster, covered by IBD segments) | 17.0442 | 14.0138 |
| d. Proportion of SNPs in 1 cM+ runs of correct phase | Mean | Median |
| Pre-Phased | 0.9368 | 0.9505 |
| IBDPhase | 0.9864 | 0.9901 |
| IBDPhase (largest supercluster) | 0.9864 | 0.9901 |
| IBDphase (SNPs covered by IBD segments) | 0.9877 | 0.9904 |
| IBDphase (largest supercluster, covered by IBD segments) | 0.9921 | 0.9926 |

**Table S2:**
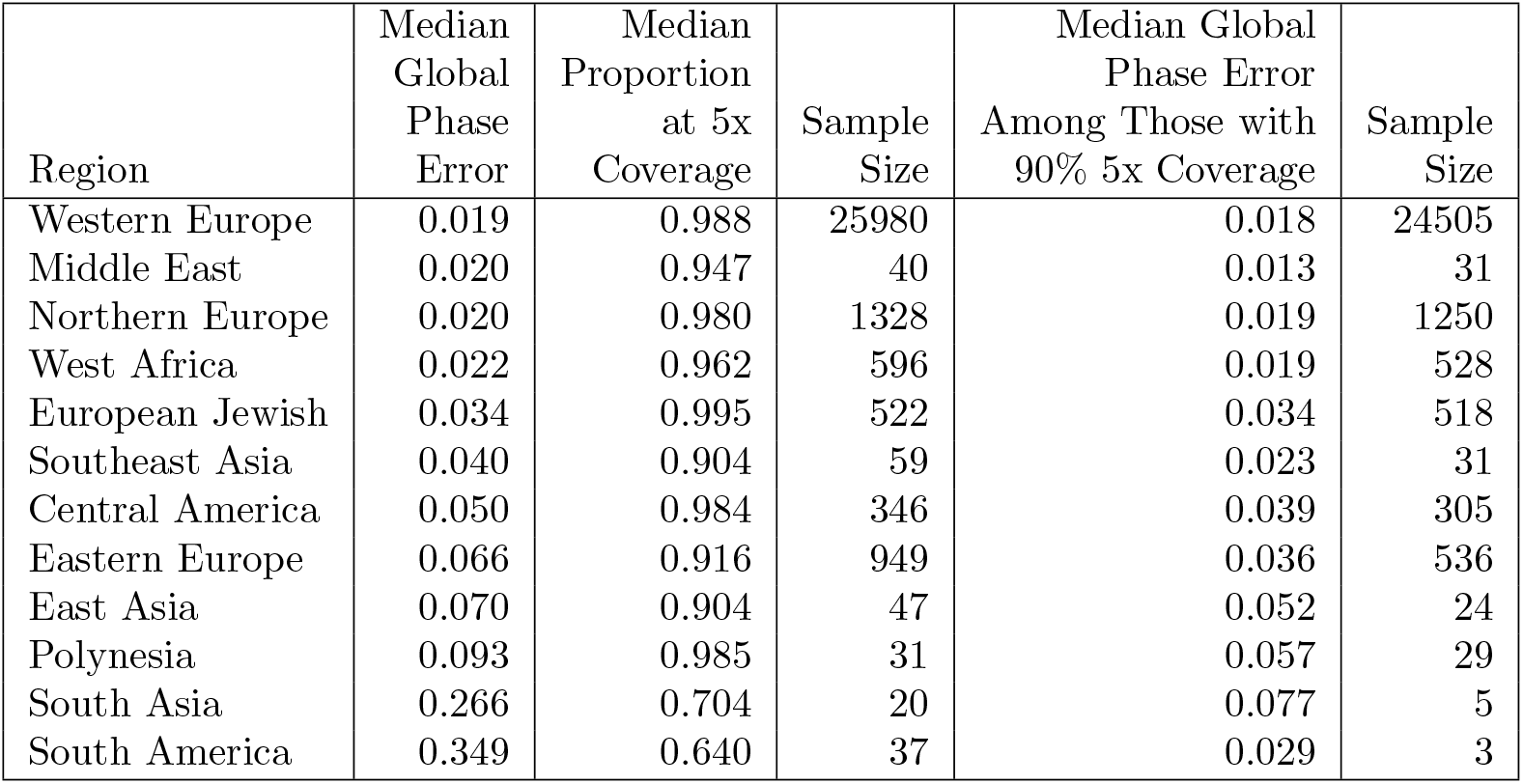
Performance Across Various Database Demographics. Each individual is classified as whichever group AncestryDNA estimates explains the origin of more DNA than any other. Global phase error is shown with all such individuals, and after limiting to those with 90% of the genome overlapped by 5 or more IBD segments.

| Region | Median Global Phase Error | Median Proportion at 5x Coverage | Sample Size | Median Global Phase Error Among Those with 90% 5x Coverage | Sample Size |
| --- | --- | --- | --- | --- | --- |
| Western Europe | 0.019 | 0.988 | 25980 | 0.018 | 24505 |
| Middle East | 0.020 | 0.947 | 40 | 0.013 | 31 |
| Northern Europe | 0.020 | 0.980 | 1328 | 0.019 | 1250 |
| West Africa | 0.022 | 0.962 | 596 | 0.019 | 528 |
| European Jewish | 0.034 | 0.995 | 522 | 0.034 | 518 |
| Southeast Asia | 0.040 | 0.904 | 59 | 0.023 | 31 |
| Central America | 0.050 | 0.984 | 346 | 0.039 | 305 |
| Eastern Europe | 0.066 | 0.916 | 949 | 0.036 | 536 |
| East Asia | 0.070 | 0.904 | 47 | 0.052 | 24 |
| Polynesia | 0.093 | 0.985 | 31 | 0.057 | 29 |
| South Asia | 0.266 | 0.704 | 20 | 0.077 | 5 |
| South America | 0.349 | 0.640 | 37 | 0.029 | 3 |

**Figure S5:**
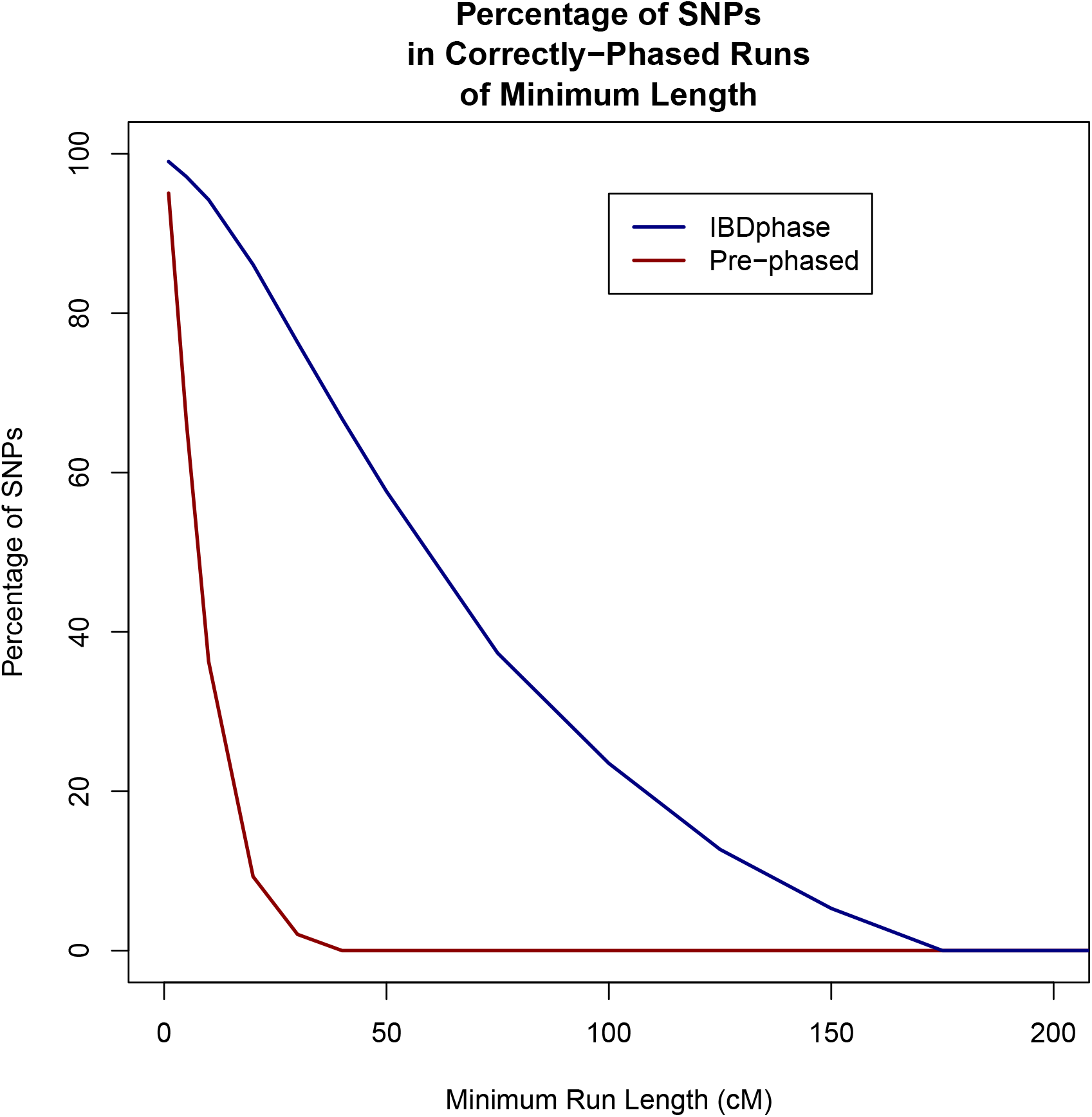
Length of Runs of Verified-Correct Phase. Distribution of the median proportion of SNPs that are part of runs of various minimum length that are in 100% agreement with the trio-phased standard. For example, a majority of SNPs phased with IBDphase are part of runs of at least 50 centimorgans that are identical to trio phase.

**Figure S6:**
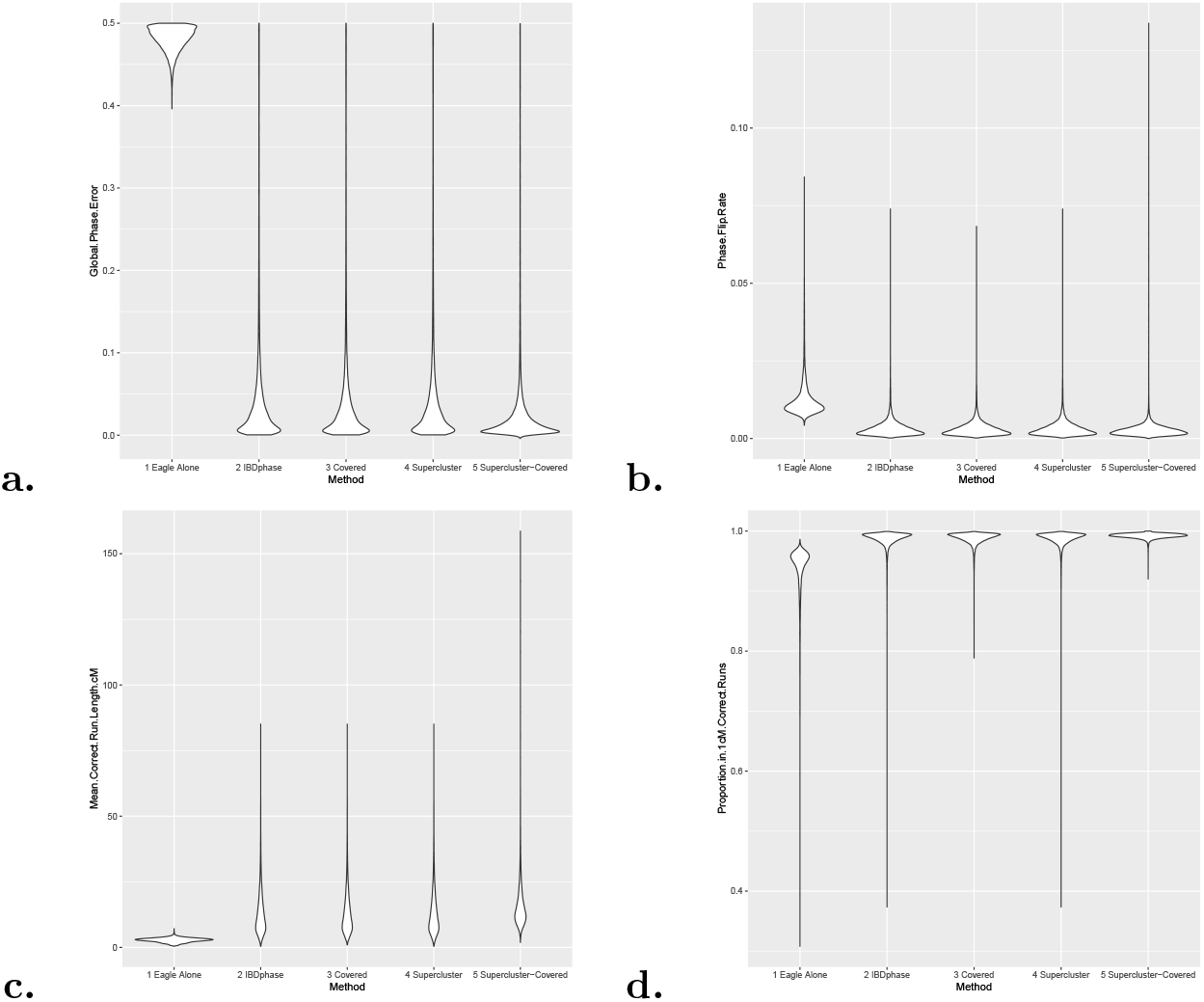
Distribution of phase error. Violin plots showing the distribution of phase error rates measured in various ways on (1) data phased with Eagle alone, (2) data phased with Eagle, then IBDPhase, (3) IBDPhase restricted to SNPs that are covered by at least one IBD segment, (4) IBDPhase restricted to sections of the genome that overlap the largest supercluster, (5) IBDPhase restricted to the largest supercluster and are covered by an IBD segment. **a**. Distribution of global (genome-wide) phase error: assuming we pair each haplotype with whichever parent data is the most favorable, how many alleles in heterozygous sites are assigned to the wrong parent? We calculate this by comparing the phase with the parents’ genotypes at sites where the correct phase is unambiguous (*i*.*e*., at least one parent is homozygous). **b**. Distribution of phase-flip rate: what proportion of (unambiguously phased) heterozygous sites are out-of-phase with the previous heterozygous site? **c**. Distribution of the average length of runs of correct phase: what is the mean length (in centimorgans) of a section of the genome that is uninterrupted by a phase flip? **d**. Distribution of the proportion of the genome that is in a run of correct phase that is at least 1 cM long: what proportion of (unambiguously phased) SNPs are in runs of at least 1 cM that are uninterrupted by a phase flip error?

**Figure S7:**
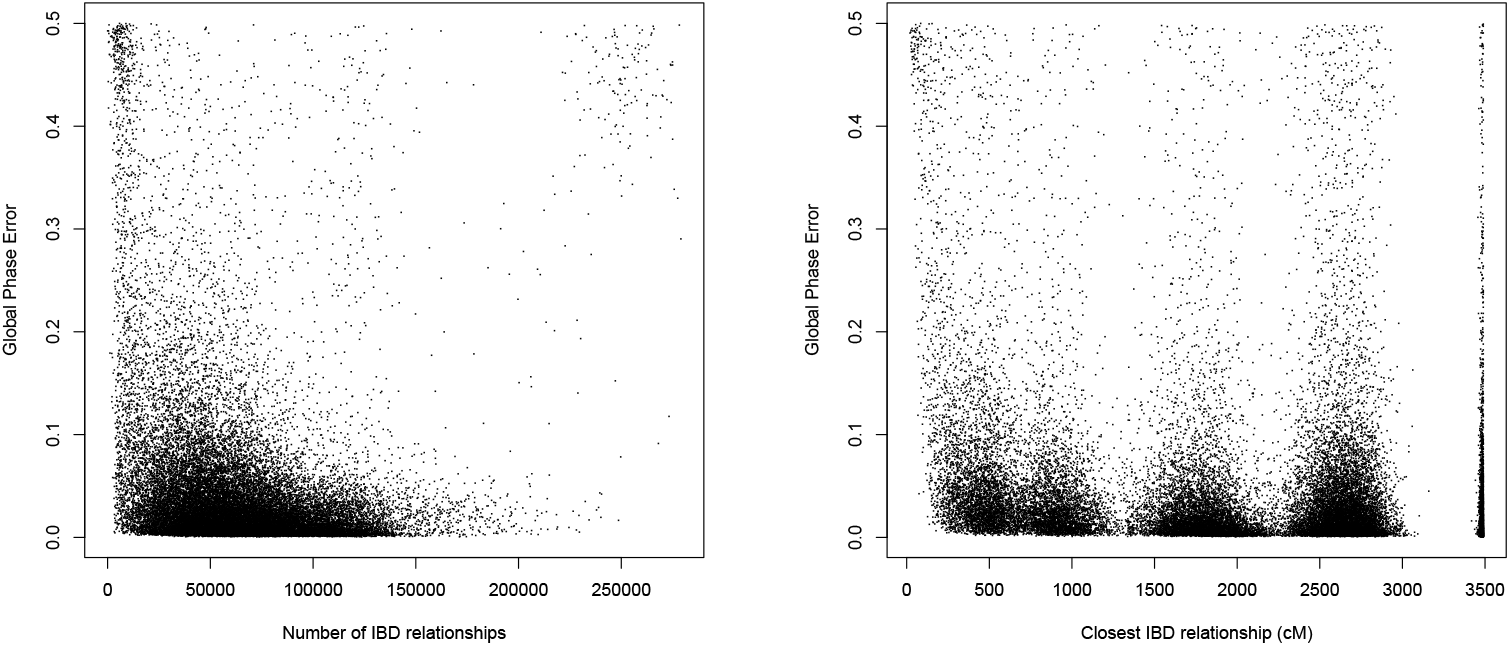
Scatterplots comparing IBD statistics and global phase accuracy. There is no strong relationship between the number of IBD segments nor the closest genetic relationship, and the accuracy of phase.

**Figure S8:**
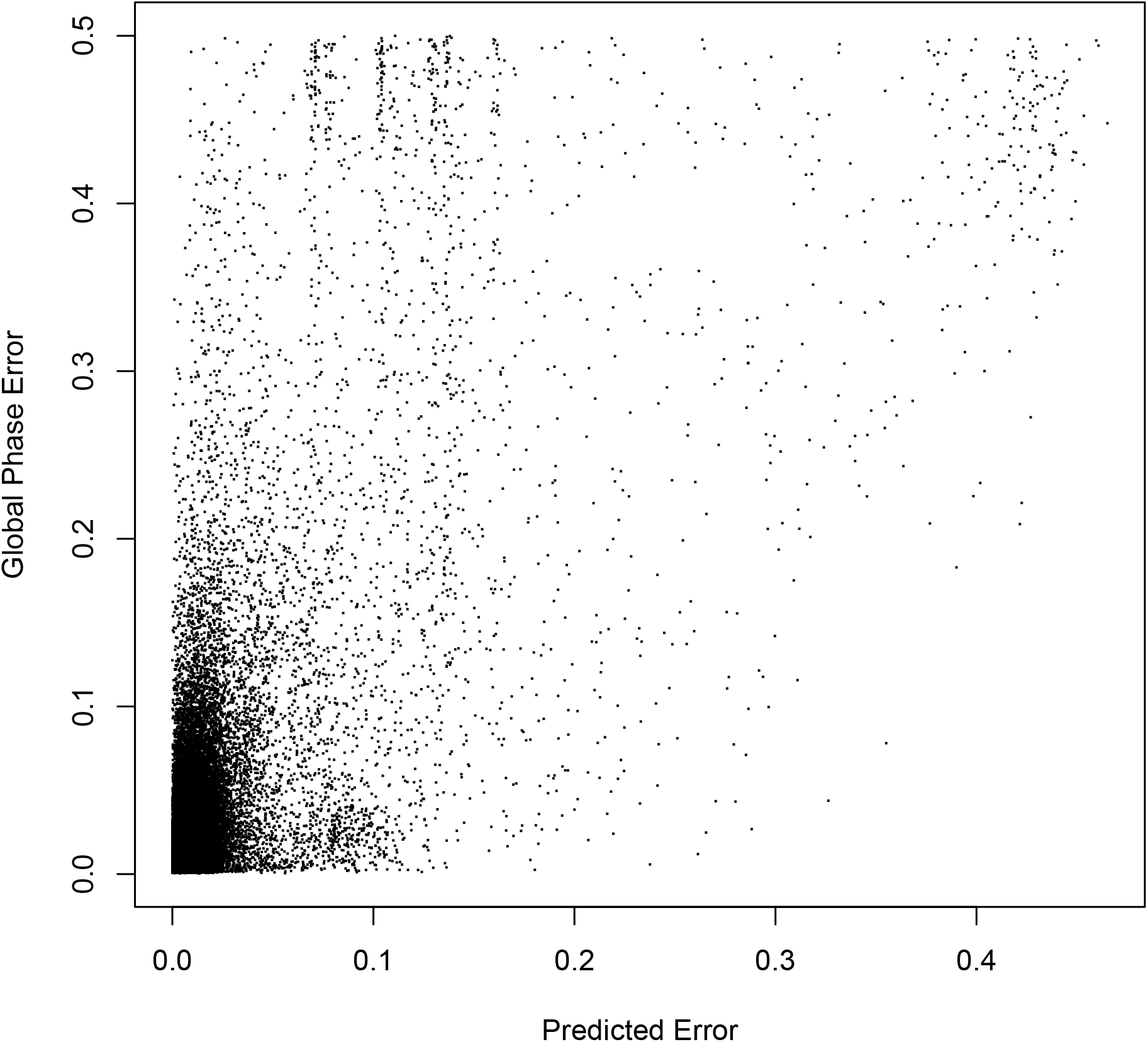
Scatterplot comparing predicted IBD clustering error and global phase accuracy.

**Figure S9:**
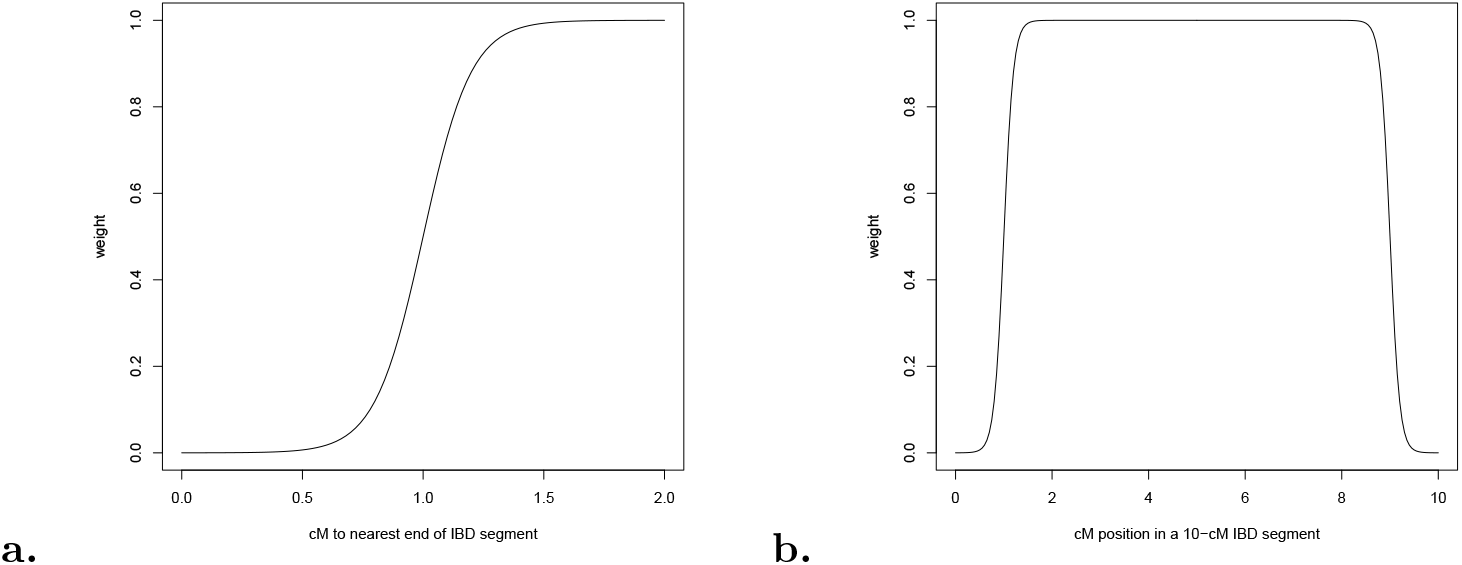
The weight function that we overlay on IBD segments when making phasing decisions based on the genotypes of multiple IBD segments. The function is *weight* = (1 + *e*^*−* 10*×*(*cM−* 1)^)^*−* 1^ where *cM* represents the distance in centimorgans to the nearest end of the IBD segment. The motivation for doing this is that it is difficult to pinpoint where DNA sharing because of IBD begins and ends, and therefore the information nearest the ends is less reliable.

**Figure S10:**
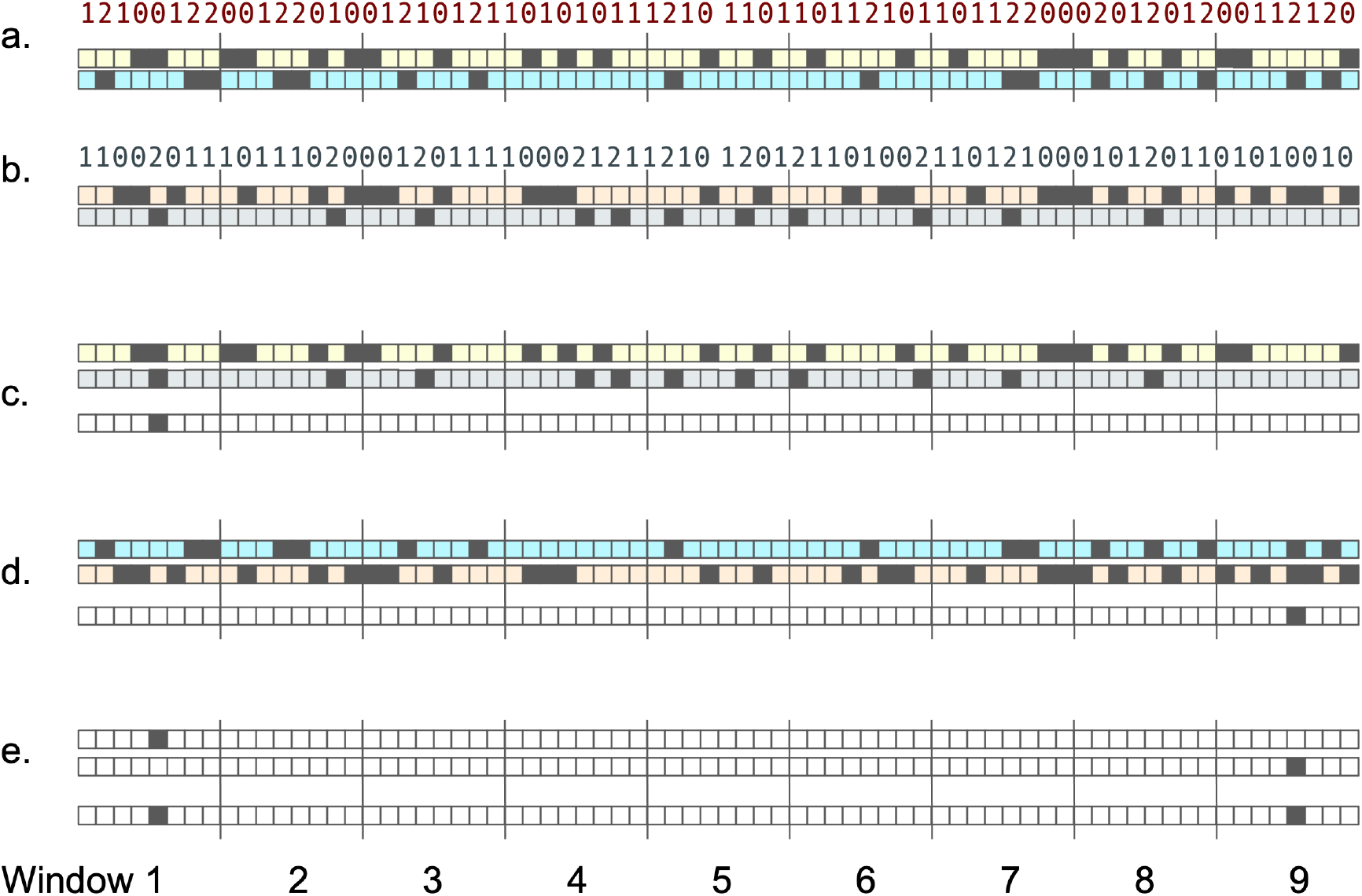
Illustration of the bitwise arithmetic used to discover IBD segments. Part **a**. shows the genotype of one individual, *A*, expressed as the number of copies of an alternate allele, 0, 1, or 2, and two corresponding bitmaps: the positions where the individual is homozygous for the reference allele, and the positions where the individual is homozygous for the alternate allele. **b**. shows the same for another individual, *B*. **c**. shows the result of applying bitwise *and* to individual *A*’s homozygous-reference bitmap and *B*’s homozygous-alternate bitmap. **d**. shows the result of applying bitwise *and* to individual *A*’s homozygous-alternate bitmap and *B*’s homozygous-reference bitmap. **e**. shows the result of applying bitwise *or* to the results from **c**. and **d**. The result in **e**. tells us that individuals *A* and *B* do not share an allele for one of the SNPs in window 1 because that window evaluates to nonzero, but there are no such homozygous opposites in windows 2-8. If that segment is long enough, we can compute the exact coordinates of the IBD segment. (The illustration uses 8-bit/8-SNP windows, but in practice we use 64-bit arithmetic.)

**Figure S11:**
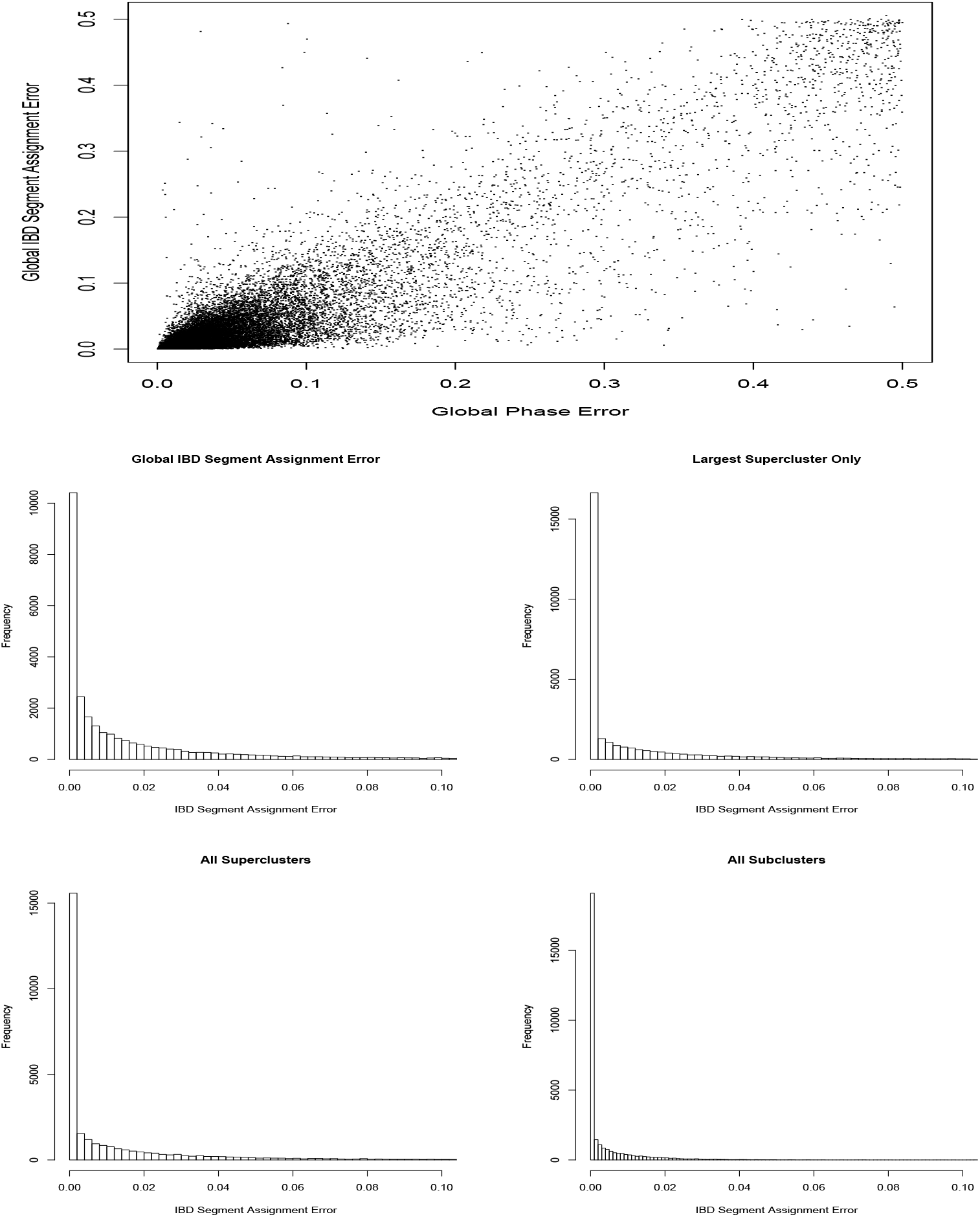
IBD Segment Assignment Error. These plots show the accuracy of assigning the IBD segments of a test set trio child to two parents, where accuracy is determined by whether the same IBD segments are also found in the IBD data of the parents. The first plot compares this measure of accuracy to global phase accuracy. The others show histograms of error distribution if we include all IBD segments, just the ones in the largest supercluster of each test set example, if we evaluate each supercluster independently of the others, and if we evaluate each subcluster independently of the others.

